# A high-quality reference genome and comparative genomics of the widely-farmed banded cricket (*Gryllodes sigillatus*) identifies selective breeding targets

**DOI:** 10.1101/2025.01.09.631893

**Authors:** Shangzhe Zhang, Kristin R. Duffield, Bert Foquet, Jose L. Ramirez, Ben M. Sadd, Scott K. Sakaluk, John Hunt, Nathan W. Bailey

**Affiliations:** School of Biology, University of St Andrews, St Andrews, Fife KY16 9TH, UK; USDA-ARS, Geospatial and Environmental Epidemiology Research Unit, Mississippi State, MS, USA; USDA-ARS, National Center for Agricultural Utilization Research, Crop BioProtection Research Unit, Peoria, IL, USA; School of Biological Sciences, Illinois State University, Normal, IL 61790, USA; McGuire Center for Lepidoptera and Biodiversity, Florida Museum of Natural History, University of Florida, Gainesville, FL, USA; School of Science, Western Sydney University, Hawkesbury Campus, Penrith NSW 2751, Australia

**Keywords:** banded cricket, cricket farming, genome-assisted breeding, *Gryllodes sigillatus*, insect protein

## Abstract

Farmed insects have gained attention as an alternative, sustainable source of protein with a lower carbon footprint than traditional livestock. We present a high-quality reference genome for one of the most commonly farmed insects, the banded cricket *Gryllodes sigillatus*. In addition to its agricultural importance, *G. sigillatus* is also a model in behavioural and evolutionary ecology research on reproduction and mating systems. We report comparative genomic analyses that clarify the banded cricket’s evolutionary history, identify gene family expansions and contractions unique to this lineage, associate these with agriculturally important traits, and identify targets for genome-assisted breeding efforts. The high-quality *G. sigillatus* genome assembly plus accompanying comparative genomic analyses serve as foundational resources for both applied and basic research on insect farming and behavioural biology, enabling researchers to pinpoint trait-associated genetic variants, unravel functional pathways governing those phenotypes, and accelerate selective breeding efforts to increase the efficacy of large-scale insect farming operations.

## Introduction

Global demand for sustainable protein sources has driven interest in alternative agriculture. Such efforts include insect farming, which can meet nutritional needs while significantly reducing environmental impacts (Grabowski et al., 2022; van Huis & Oonincx, 2017). Insects are known for their favourable feed conversion efficiency. They require significantly less land, water, and energy resources compared to traditional livestock, exploit diverse dietary proteins (Morales-Ramos et al., 2020; Kasdorf et al., 2025) and produce fewer greenhouse gases (Lange & Nakamura, 2023). These advantages make them an environmentally friendly option for addressing food security challenges (Huis et al., 2021; van Huis & Oonincx, 2017). Crickets (suborder Ensifera) have been farmed and consumed by humans for millennia, and over 60 species are known to be consumed across 49 countries (Magara et al., 2021). Among these, the banded cricket (*Gryllodes sigillatus*, Figure 1a) has gained considerable attention due to its rapid growth rate, high reproductive capacity, and palatability, making it a preferred species for large-scale farming operations (Kong et al., 2024; Magara et al., 2021).

**Figure 1.**
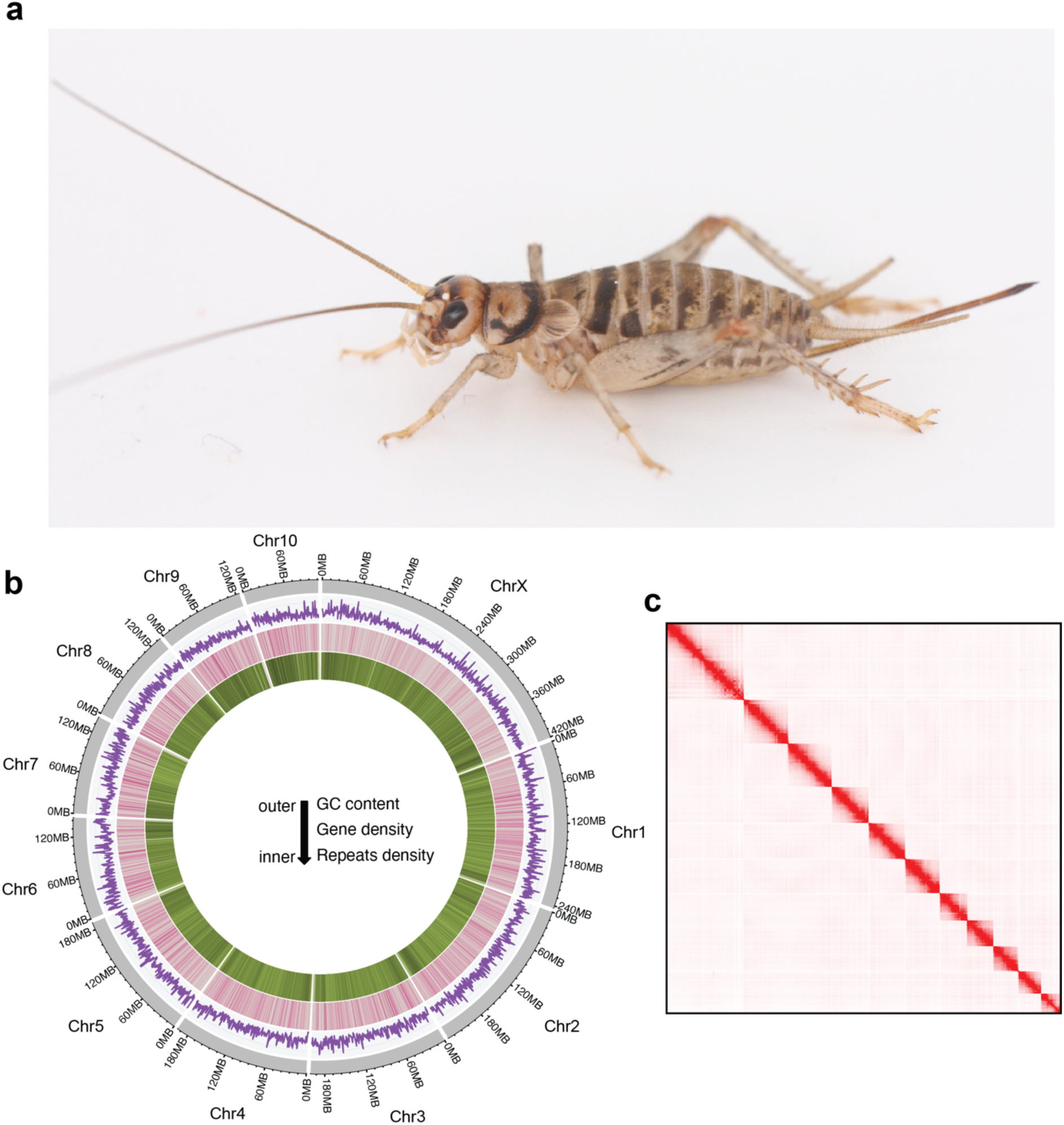
*Gryllodes sigillatus* and its genome characteristics. **(a)** A female *G. sigillatus*. **(b)** Features of the *G. sigillatus* genome. Outer track (grey) illustrates the 11 chromosomes, with the largest identified as the putative X chromosome on the basis of a published karyotype study (You et al., 2007). Genome position is indicated by the outer numbers. The inner purple track shows GC content, and the inner pink and green tracks show genome-wide gene and repeat density, respectively. **(c)** Hi-C contact map of the genome assembly indicates high contiguity and coverage. **Photo credit**: NW Bailey.

Despite the growing commercial significance of *G. sigillatus*, publicly available genetic resources for this species are limited. This scarcity of genetic information constrains the potential for selective breeding programs that could enhance desirable traits such as nutritional value, growth rate, and reproductive efficiency. Recent research on *G. sigillatus* has identified key environmental factors that influence such traits, including temperature, food stress (Rapkin et al., 2018; Kasdorf et al., 2025; Muzzatti et al., 2024; Muzzatti et al., 2025a), mating strategies (Sakaluk et al., 2019) and pests (Muzzatti et al., 2025b). However, genome-based selection has become central to modern breeding in a wide variety of agricultural species, resulting in rapid and efficient improvement of specific traits (Johnsson, 2023). Beyond their economic importance, field crickets – including *G. sigillatus* - have long been a study organism in evolutionary and behavioural ecology (Bateman & MacFadyen, 1999; Champagnon & Cueva del Castillo, 2008; Sakaluk et al., 2002; Sakaluk et al., 2019). Males of this polygynandrous species produce a costly nuptial gift – the spermatophylax – which they transfer to females upon mating, making *G. sigillatus* a prominent research model for examining mating system evolution, mate choice, and sexual selection (Ivy & Sakaluk, 2005; Sakaluk, 1984, 1991). A high-quality genome assembly serves as a foundational resource for such efforts, enabling researchers to pinpoint trait-associated genetic variants, unravel functional pathways governing those phenotypes, and accelerate both selective breeding efforts and fundamental research in behavioural and evolutionary biology (Sinha et al., 2021).

In this study, we produced a high-quality, annotated, chromosome-level reference genome for *G. sigillatus* by integrating long-read PacBio HiFi data with high-throughput chromosome conformation capture (Hi-C) data. Using comparative genomic approaches, we refined the phylogenetic placement and evolutionary history of *G. sigillatus*. We identified candidate gene families linked to economically significant traits that underwent changes during the divergence of *G. sigillatus* from sister taxa and illuminate gene family expansions and contractions relevant to the unique biology of this species. This research provides a foundational resource for the genetic analysis and farming of *G. sigillatus*, offering valuable insights into its genomic features and agriculturally important traits, as well as for basic research on the evolution of mating systems and reproductive behaviour.

## Methods

### Sampling and DNA sequencing

For whole genome sequencing, an individual *Gryllodes sigillatus* female was taken from an inbred laboratory stock line descended from approximately 500 adults collected at Las Cruces, New Mexico, in 2001 (Ivy & Sakaluk, 2005). We extracted high molecular weight genomic DNA from liquid nitrogen flash-frozen cricket legs. Following an initial wash with 1× PBS buffer, DNA was extracted using the Macherey-Nagel™ NucleoBond™ High Molecular Weight DNA Kit. DNA quality and concentration were assessed with both Nanodrop and Qubit. Sequencing library preparation and whole genome sequencing were performed using the PacBio SMRT^®^ system by BGI Genomics, resulting in 96.25 Gb of HiFi reads with a coverage depth of approximately 48×. Additionally, flash-frozen muscle tissue was used to prepare and sequence a Hi-C library, also by BGI Genomics, resulting in 151.48 Gb of ca. 72× Hi-C reads (Table S1).

### Bulk RNA-seq

RNA-seq data used to facilitate genome annotation was generated using two different methods. This was done so as to include reads from as wide a range of tissues as possible to increase the completeness of the genome annotation. We used data from whole bodies and accessory glands of males, plus whole bodies, ovaries, and heads of females (Table S2).

Whole body RNA-seq data was generated from laboratory stock crickets as follows. These were descended from 500 adult *G. sigillatus* collected in Las Cruces, New Mexico (USA) in 2001 (Ivy & Sakaluk, 2005). Rearing followed previously described methods (Duffield et al., 2021, 2022). Approximately 500 crickets were housed in ventilated, 55 L plastic storage bins packed with egg carton. Crickets were given *ad libitum* food (approximately equal parts Purina Cat Chow Complete Cat Food and Teklad Global 18% Protein Extruded Rodent Diet pellets) and water (glass vials plugged with moist cotton). Colonies were housed in an environmental chamber at 32 °C on a 16:8 hour light:dark cycle. Experimental crickets were frozen and stored at -80 °C at 7 days post-eclosion and shipped on dry ice to LC Sciences (Houston, TX) for further processing. Total RNA was extracted using Trizol (Invitrogen, USA) following the manufacturer’s procedure. Total RNA quality and quantity were analysed using a Bioanalyzer 2100 and RNA 6000 Nano LabChip Kit (Agilent, USA), with samples obtaining RIN number >7.0. Then, mRNA was purified in two rounds using Dynabeads Oligo (dT) (Thermo Fisher, USA) and subsequently fragmented with a Magnesium RNA Fragmentation Module (New England Biolabs, USA) at 94℃ for 5-7min. The cleaved RNA fragments were reverse-transcribed to cDNA using SuperScript™ II Reverse Transcriptase (Invitrogen, USA) and then used to synthesize U-labelled second-stranded DNAs with *E. coli* DNA polymerase I (New England Biolabs, USA), RNase H (New England Biolabs, USA), and dUTP Solution (Thermo Fisher, USA). Dual-index adapters were ligated to the fragments, and size selection was performed with AMPureXP beads. After the heat-labile UDG enzyme (New England Biolabs, USA) treatment of the U-labelled second-stranded DNAs, the ligated products were amplified with PCR by under the following conditions: initial denaturation at 95℃ for 3 min; 8 cycles of denaturation at 98℃ for 15 sec, annealing at 60℃ for 15 sec, and extension at 72℃ for 30 sec; and then final extension at 72℃ for 5 min. The average insert size for the final cDNA libraries was 300±50 bp. Sequencing was performed using 2×150bp paired-end sequencing (PE150) on an Illumina Novaseq™ 6000 following the vendor’s recommended protocol (Table S2).

RNA-seq data for male accessory glands, female ovaries, and female heads were generated from a lab-maintained line (Burns-Dunn et al., 2024; McKermitt et al., 2024), that originated from the same crickets collected in Las Cruces, 2001 (Ivy & Sakaluk, 2005). Approximately 300 adults were housed in ventilated 62.5 L plastic storage bins packed with egg carton. Nymphs were maintained in 6 L bins. Crickets were given *ad libitum* food (approximately equal parts Purina Cat Chow Complete Cat Food and Teklad Global 18% Protein Extruded Rodent Diet pellets) and water (glass vials plugged with moist cotton). Colonies were maintained at a constant temperature (30 ± 2 °C) and under a reversed photoperiod (14:10 hr light:dark cycle). Seven-day old crickets were mated, and two hours later they were stunned on ice for tissue dissections. Each tissue was immediately snap frozen in liquid nitrogen and stored -80 °C until further processing. RNA extractions were performed as described in Foquet et al. (2023). RNA was extracted using a Trizol (Thermo Fisher Scientific) and 1-bromo-3- chloropropane (BCP, Acros Organics) phase separation using standard procedures, followed by a DNAse treatment using a TURBO DNA-free kit (Thermo Fisher Scientific). RNA qualities and concentration were measured on a MultiSkan GO microplate spectrophotometer with a μDrop adapter plate (ThermoScientific). Only RNA with final 260/230 and 260/280 values higher than two were used for RNA sequencing. RNA samples were sent to Novogene for library preparation, sequencing, and read pre-processing. Sample quality was assessed with a Bioanalyzer 2100 (Agilent, USA), with samples obtaining RIN number >5.0 used for sequencing. RNA was processed with an Illumina TruSeq Stranded mRNA Kit with polyA enrichment, and sequencing of paired end 150bp reads on a NovaSeq were performed by Novogene.

### Genome assembly

We used HiFiasm v0.19.5-r587 (Cheng et al., 2021), with default settings to generate a draft assembly of *G. sigillatus*. The primary assembly was retained for subsequent refinement and analysis. To eliminate duplicates within the contig-level assembly, Purge_Dups v1.2.6 (Guan et al., 2020) was employed. This identified redundant contigs by mapping the original HiFi reads back to the assembly, removing repetitive or duplicated sequences that could inflate assembly size and misrepresent genomic content. To remove mitochondrial sequences from the draft assembly, MitoHiFi v3.2.1 (Uliano-Silva et al., 2023) was used. This tool identified and removed contigs associated with the mitochondrial genome by comparing them against the published *G. sigillatus* mitochondrial sequence (MW365703.1, Yang et al., 2021), ensuring that only nuclear genomic sequences remained in the primary assembly.

Hi-C data were then incorporated to scaffold the contigs and achieve chromosome-level assembly following Arima Hi-C mapping guidelines (https://github.com/ArimaGenomics/mapping_pipeline). Briefly, this involved mapping with BWA-MEM2 v2.0pre2 (Vasimuddin et al., 2019) and removing PCR duplicates using Picard v3.1.0 (https://broadinstitute.github.io/picard). All Hi-C reads were mapped to the contig-level assembly, providing spatial genomic interaction information. YahS v1.1a-r3 (Zhou et al., 2023) was used to scaffold contigs based on these mapping results, integrating long-range contact information to order and orient contigs accurately. To further refine scaffolding, we used Juicer tools (https://github.com/aidenlab/Juicebox) and Juicebox v3.0.0 (https://github.com/aidenlab/JuicerTools) to visualise the Hi-C contact heatmap and then manually inspect and correct misassembled regions.

Once the final scaffolds were generated, BlobTools2 (Challis et al., 2020) was used to screen for potential contaminant sequences originating from other species. This step identified 23 scaffolds that likely represented contamination, and these were removed to produce the final, cleaned scaffold-level genome assembly. This comprehensive, iterative workflow maximised the likelihood of obtaining an accurate, high-quality chromosome-level *G. sigillatus* genome assembly suitable for further genomic analyses and annotation.

### Genome quality assessment

Genome assembly statistics, including total length, N50, and GC content, were calculated using SeqKit v2.4.0 (Shen et al., 2016) to provide a comprehensive overview of the assembly quality.

To assess the completeness of the assembled genome, BUSCO v5.4.6 (Manni et al., 2021) was employed, aligning the nucleotide sequences against the Arthropoda dataset (Odb10). This analysis enabled the evaluation of the assembly’s coverage and accuracy based on the presence of conserved single-copy orthologs representative of the Arthropoda lineage.

To evaluate the integrity and accuracy of the genome assembly, we employed the k-mer-based method Merqury v1.3 (Rhie et al., 2020). This approach involves mapping k-mers derived from raw HiFi reads to the contig-level assembly, enabling a detailed assessment of assembly quality. Specifically, Merqury calculates the Quality Value (QV) score, which serves as a metric for the overall accuracy of the assembly and estimates the error rate at the level of individual bases.

For evaluating the completeness of the protein-coding gene annotation (see below), BUSCO was again utilised with the Insecta dataset in ‘transcriptome’ mode, which is designed to analyse transcriptome assemblies. This allowed for a thorough assessment of gene prediction quality, ensuring that the annotated protein-coding genes were representative and complete. By using both genome and transcriptome completeness assessments, we ensured a robust evaluation of the assembly’s accuracy and the integrity of its functional annotations.

### Genome annotation

To identify repetitive elements in the genome sequence, three sequential methods were employed to ensure thorough coverage. First, Tandem Repeats Finder v4.09 (TRF; Benson, 1999) was used to detect simple sequence repeats. The repetitive regions identified by TRF were masked with ‘N’ in the genome sequence, effectively obscuring these regions for downstream analyses. Next, transposable elements (TEs) within the genome were identified using RepeatMasker v4.1.2-p1 (Tarailo-Graovac & Chen, 2009), with the previously masked genome sequence serving as input. During this step, the genomic sequences were aligned against the Repbase database v20181026 (Bao et al., 2015) to specifically search for TEs homologous to the Arthropoda lineage. The genome was again subjected to hard-masking based on the identified TEs. Lastly, RepeatModeler2 (Flynn et al., 2020) was applied to conduct a self-alignment of the genomic sequences to uncover any additional repetitive elements. The results from RepeatModeler2 were incorporated into a custom repeat database, and RepeatMasker was run once more to identify repeats based on alignment hits. All repetitive elements identified through this comprehensive protocol were soft-masked in subsequent analyses, preserving their sequence while distinguishing them for further study.

For gene prediction, we integrated both homologous protein sequences and RNA-seq data. Protein sequences from *D. melanogaster* (BDGP6.46, Ensembl; Harrison et al., 2024), *G. bimaculatus* (Ylla et al., 2021), *T. occipitalis* (Kataoka et al., 2020) and *T. oceanicus* v2 (Zhang et al., 2024), and reviewed orthologs from Arthropoda were obtained from SwissProt to guide the annotation process. Additionally, RNA-seq data from multiple tissues and both sexes were utilised to facilitate transcriptome-based annotation. Prior to gene prediction, raw RNA-seq reads were quality-trimmed using Fastp v0.23.2 (Chen et al., 2018) and subsequently mapped to the reference genome with HISAT2 v2.2.0 (Kim et al., 2015), ensuring accurate alignment for transcript assembly.

To construct the final gene set, we employed a combined annotation strategy that integrated results from three distinct gene prediction approaches. First, BRAKER3 v3.0 (Gabriel et al., 2024) was used for *ab initio* gene prediction, leveraging both protein and transcript evidence to train and predict gene structures. Second, Miniprot v0.10-r225 (H. Li, 2023) was employed to map homologous protein sequences from closely related species and Arthropoda orthologs to the reference genome, providing potential coordinates and open reading frame (ORF) structures of protein-coding genes. Third, RNA-seq data were assembled using two different tools, StringTie v2.2.1 (Shumate et al., 2022) and Trinity v2.14.0 (Grabherr et al., 2011), to capture transcript diversity. The Trinity-assembled transcriptome was filtered to retain sequences longer than 1 Kb, and redundancy was reduced using CD-HIT v4.8.1 (W. Li & Godzik, 2006). The PASA pipeline v2.5.3 (https://github.com/PASApipeline/PASApipeline) was then utilised to predict gene structures by mapping these transcripts to the reference genome. For the transcriptome assembled by StringTie, potential ORFs were identified using TransDecoder v5.7.1 (https://github.com/TransDecoder/TransDecoder), and coordinates were mapped to the reference genome. Finally, the EvidenceModeler v2.1.0 (Haas et al., 2008) was employed to integrate all prediction results into a unified gene set, while the PASA pipeline refined the final gene models by aligning the transcriptome to predicted gene structures. This comprehensive approach resulted in a high-confidence, annotated gene set for the target genome.

### Functional annotation

The functions of protein-coding genes were annotated by mapping the isoform sequences against multiple publicly available functional databases. For Non-redundant (Nr; Sayers et al., 2022) and Swiss-Prot/TrEMBL (Boeckmann et al., 2003), protein sequences were mapped to an Insecta-specific subset using DIAMOND blastp v2.0.14.152 (Buchfink et al., 2021), enabling accurate identification of protein functions within this taxonomic group. To identify and annotate functional domains within the protein sequences, InterProScan (Paysan-Lafosse et al., 2023) was employed, which integrates multiple signature databases for a thorough domain analysis. Additionally, EggNOG mapper (Cantalapiedra et al., 2021) was utilised to conduct EggNOG (Huerta-Cepas et al., 2019) analysis, from which Gene Ontology (GO) terms were extracted to provide insights into the biological processes, molecular functions, and cellular components associated with each protein. Finally, KEGG pathway (Kanehisa et al., 2016) annotations were acquired using the KAAS online server (Moriya et al., 2007), allowing for the identification of biochemical pathways and the functional context of the proteins within broader metabolic networks.

### Ortholog group identification and phylogenetic analysis

Orthologous protein sequences from six other genomes, including *Anabrus simplex* (NCBI RefSeq accession: GCF_040414725.1 by United States Department of Agriculture), *Gryllus longicercus* (Szrajer et al., 2024), *Gryllus bimaculatus* (Ylla et al., 2021), *Teleogryllus oceanicus* (Zhang et al., 2024), *Teleogryllus occipitalis* (Kataoka et al., 2020) and *Schistocerca americana* (iqSchAmer2.1, https://www.dnazoo.org/assemblies/schistocerca_americana), along with our *G. sigillatus* proteins, were collected for ortholog group identification using OrthoFinder v2.2.5 (Emms & Kelly, 2019). To construct a robust phylogenetic framework, we focused on single-copy orthologs, which were used to build a maximum-likelihood phylogenetic tree, providing insights into the evolutionary relationships among these species. To estimate divergence times, we incorporated reference divergence times from fossil evidence obtained from the TimeTree database (Kumar et al., 2022), ensuring that our analysis was anchored to well-established evolutionary benchmarks. The Bayesian inference tool MCMCTree, part of the PAML v4.10.7 package (Reis & Yang, 2011), was then employed to estimate divergence times across all nodes within the phylogeny using only four-fold sites within all single-copy gene sequences.

### Gene family analysis

Using the ultrametric tree generated by MCMCTree, we analysed the expanded and contracted gene families across all nodes of the phylogeny with CAFE5 v1.1 (Mendes et al., 2021), a software designed for modelling gene family evolution. To optimise the analysis, we tested a range of discrete gamma rate (*k*) values from 2 to 6, which account for the variation in evolutionary rates across gene families. The *k*=4 that yielded the highest maximum likelihood was selected for the final model, ensuring an accurate fit to the data. To identify gene families that experienced significant changes specifically in *G. sigillatus*, we conducted a Likelihood Ratio Test (LRT), which enabled us to distinguish gene families that had undergone statistically significant expansions or contractions.

### Statistical analysis and visualisation

Statistical analyses were performed using R v4.0.2. Diagrams were generated using ggplot2 in the tidyverse package (Wickham et al., 2019). The genomic circos plot was generated using shinyCircos v2.0 (Wang et al., 2023).

## Results and Discussion

### Chromosome-level genome assembly of *G. sigillatus*

We generated a high-quality, chromosome level annotated reference genome for *G. sigillatus* (Figure 1b; Table 1; Table S3). The completed assembly has a total size of 2.17 Gb and comprises 11 pseudochromosomes (Figure 1c; Table S4), consistent with the known karyotype of *G. sigillatus* (You et al., 2007). Based on the latter karyotype study, the longest scaffold in the assembly was marked as chromosome X and autosomes were named by order of length. The assembly has a scaffold N50 of 200.35 Mb and a maximum scaffold length of 420.78 Mb (Figure 1b, Table S3), and k-mer based quality estimation yielded a QV score of 61.21 and per-base error rate of 7.56×10^-7^ (Table S4), indicating a highly contiguous and complete genome assembly. To further assess the completeness and quality of the assembly, we performed Benchmarking Universal Single-Copy Orthologs (BUSCO) analysis using the insecta_odb10, which yielded a BUSCO recovery score of 99.49% (Table S5).

**Table 1.**
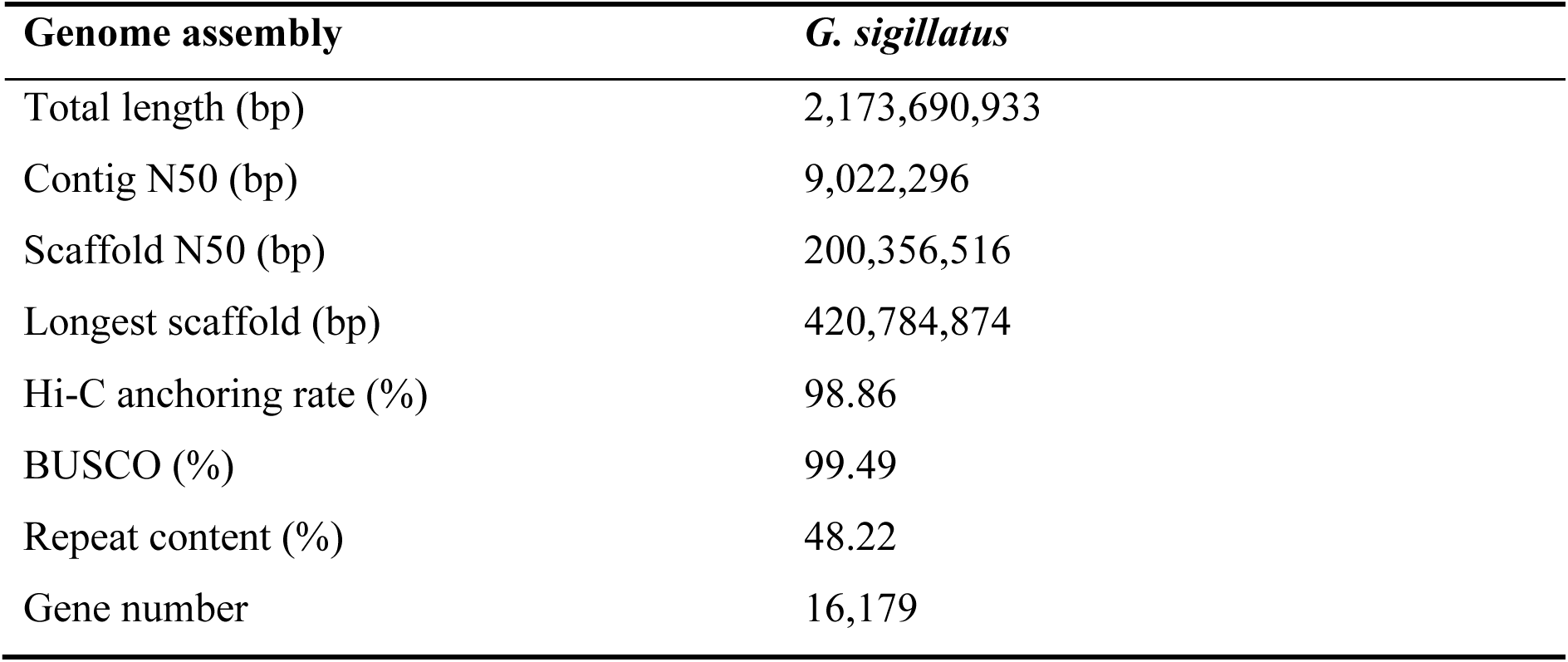
Summary of *Gryllodes sigillatus* genome assembly and annotation.

| Genome assembly | *G. sigillatus* |
| --- | --- |
| Total length (bp) | 2,173,690,933 |
| Contig N50 (bp) | 9,022,296 |
| Scaffold N50 (bp) | 200,356,516 |
| Longest scaffold (bp) | 420,784,874 |
| Hi-C anchoring rate (%) | 98.86 |
| BUSCO (%) | 99.49 |
| Repeat content (%) | 48.22 |
| Gene number | 16,179 |

Genome annotation revealed that repetitive sequences accounted for 48.22% of the genome, including DNA transposons (16.22%), long interspersed nuclear elements (LINEs, 24.13%), short interspersed nuclear elements (SINEs, 1.81%), and long terminal repeat (LTR) retrotransposons (0.07%) (Figure 1b; Table S6). This repeat content is comparable to that observed in related species (Szrajer et al., 2024; Ylla et al., 2021; Zhang et al., 2024), suggesting relatively conserved genomic architecture and evolutionary patterns of transposable elements. We predicted a total of 16,179 protein-coding genes using a combination of *ab initio* prediction and evidence-based methods, incorporating RNA-seq data (Figure 1b; Table S2) and protein homology. Functional annotation was successful for 84.56% of the total predicted genes, using alignment against several public databases including the Non-redundant Protein Sequence Database (Nr), TrEMBL, Gene Ontology (GO), EggNOG, and the Kyoto Encyclopedia of Genes and Genomes (KEGG).

### Genome evolution and phylogenetic analysis

Our study provides the first reference genome within the *Gryllodes* genus. We also performed a phylogenetic analysis that broadly supports the known evolutionary history of *Gryllodes* (Song et al., 2020). A total of six Ensiferan and one Caeliferan genome (the latter as an outgroup) were selected to identify 15,123 orthogroups of which 4,157 comprised single-copy orthologs (Figure S1). A maximum-likelihood species tree was constructed using concatenated alignments of these orthologs, confirming previously-reconstructed evolutionary relationships (Song et al., 2020), validating the inferred speciation timeline, and positioning *Gryllodes* within its evolutionary context (Figure 2a). Divergence time estimates indicate that *Gryllodes* diverged from other *Gryllidae* species approximately 141-199 million years ago (Mya).

**Figure 2.**
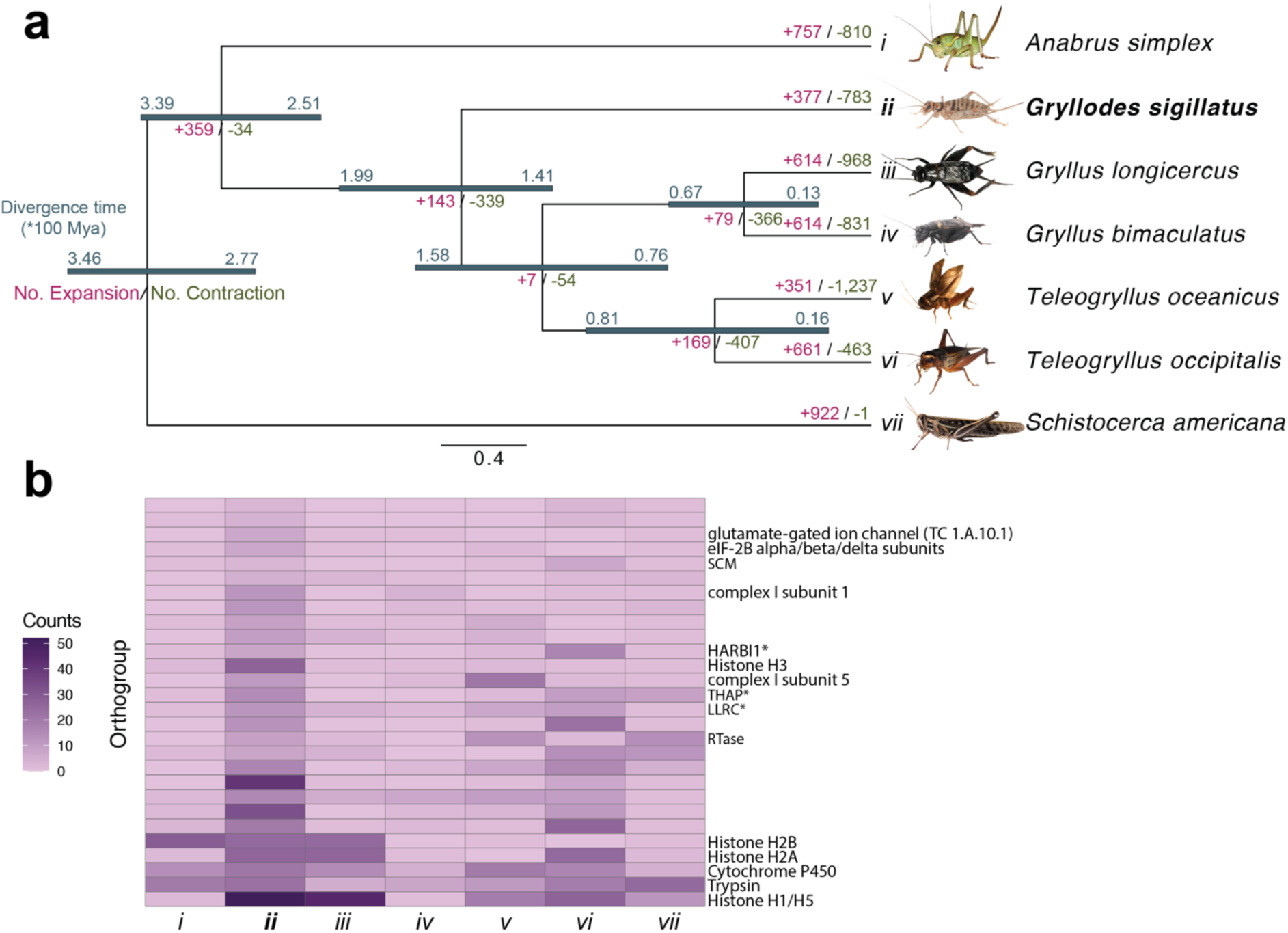
Evolution of gene family expansion and contraction in selected orthopterans. **(a)** Phylogenetic tree illustrates the divergence times and gene family evolution across seven Orthoptera species. Branch lengths represent evolutionary distances, with upper and lower divergence time estimates (in units of 100 million years ago, Mya) labelled along each branch. Numbers in red and green indicate the number of significantly expanded (+) and contracted (−) gene families along each lineage, respectively. **(b)** Heatmap illustrating counts of gene families that are significantly expanded in *G. sigillatus* (*ii*), with darker shades representing higher counts. Gene families marked with an asterisk (**\***) represent those that failed to match a verified family name across Insecta but have homologous sequences in human gene families, indicating possible highly conserved functions. **Photo credits:** *Anabrus simplex*, *Gryllodes sigillatus*, *Gryllus bimaculatus* and *Teleogryllus oceanicus* by NW Bailey; *Gryllus longicercus* by Jeff Hollenbeck, modified by Shangzhe Zhang, licensed under CC BY-ND-NC 1.0; *Teleogryllus occipitalis* by Takaaki Hattori, modified by Shangzhe Zhang, licensed under CC-BY 4.0; *Schistocerca americana* by Brandon Woo with permission granted.

Analysis of gene family evolution across orthopterans revealed 377 expanded and 783 contracted gene families in *G. sigillatus* (Figure 2a). Focusing on gene families that exhibited significant changes, we identified 28 that were notably expanded in *G. sigillatus* relative to other orthopteran species (Likelihood Ratio Test, *P* < 0.01), consisting of 146 protein-coding genes in *G. sigillatus* (Table S7). Many of these expanded families are linked to histone-related functions. Histones are central to nucleosome formation, which packages DNA within the nucleus and regulates gene expression through chromatin remodelling. In insects, histone modifications influence development (Glastad et al., 2019), stress responses (Gupta & Nair, 2024), and genome stability (Glastad et al., 2019), and the expansion of histone-related gene families in *G. sigillatus* may be associated with an enhanced capacity for chromatin remodelling and genome organisation, consistent with adaptations that enable *G. sigillatus* to be resilient in diverse ecological niches.

Other expanded gene families are associated with metabolism and energy production, including those encoding succinyl-CoA ligase (SCL), NADH dehydrogenase (complex I subunit 1), and trypsin. Succinyl-CoA ligase, a key enzyme in the tricarboxylic acid (TCA) cycle, catalyses the conversion of succinyl-CoA to succinate, producing ATP or GTP through substrate-level phosphorylation. An expansion in SCL genes may enhance TCA cycle efficiency, boosting energy production to meet increased metabolic demands for activities such as locomotion, growth, and reproduction; NADH dehydrogenase, a mitochondrial complex I component, initiates the electron transport chain by transferring electrons from NADH to ubiquinone, driving ATP synthesis via oxidative phosphorylation (Garcia et al., 2017). The expansion of genes encoding NADH dehydrogenase subunits could increase electron transport efficiency, improving ATP generation. Energy metabolism pathways and their evolution in other systems have been linked to the immunity-metabolism interface (Dolezal et al., 2019), detoxification of and resistance to xenobiotics (Gospodaryov et al., 2020), biological invasion (Wang et al., 2013), dispersal and flight (Yang et al., 2014; Treidel et al., 2023). While flight performance is less relevant in *G. sigillatus*, flight muscles are used for male reproductive signaling – acoustic advertisement song – which has been shown to be energetically constrained (Houslay et al., 2017). Overall, metabolic enhancement may further provide *G. sigillatus* with greater resilience to environmental fluctuations in energy availability, supporting sustained activity and survival across diverse conditions. Concentrating on these gene families during selective breeding efforts, for example through marker-assisted selection, could facilitate genetic improvements in nutritional requirements, energy efficiency, growth rates, and reproductive success.

The expansion of the trypsin gene family suggests adaptations in digestive processes. Trypsins are serine proteases involved in protein digestion in the insect midgut, hydrolysing peptide bonds to aid nutrient absorption (Meriño-Cabrera et al., 2022; Muhlia-Almazán et al., 2008). An increased repertoire of trypsin genes may allow *G. sigillatus* to more effectively exploit diverse dietary proteins, supporting rapid growth and high reproductive output. Trypsins are also implicated in immune responses, such as activating the prophenoloxidase system involved in melanisation and pathogen defence (González-Santoyo & Córdoba-Aguilar, 2012; Kanost & Gorman, 2008); thus, their expansion may also enhance immune capabilities in *G. sigillatus*. Determining whether such genes involving immunity are associated with increased resistance to pests and pathogens in a farming setting can help identify strains with specific haplotypes that strengthen immunity to pests, reduce mortality rates, and minimise reliance on chemical treatments. An accurate assessment of genetic variation at such loci across the global range of farmed cricket species is a priority for future work, as the evolutionary potential for responses to artificial selection are currently unknown, and natural variation is likely to be considerably higher than that available in farmed stock populations.

Finally, the expansion of the cytochrome P450 (CYP) gene family, known for its roles in detoxification and the immune response, was observed (Feyereisen, 2012). Cytochrome P450 enzymes are involved in the metabolism of endogenous substrates and detoxification of xenobiotics (Nauen et al., 2022), including plant secondary metabolites and synthetic insecticides (Dermauw et al., 2020). Additionally, CYP enzymes contribute to the synthesis and degradation of hormones such as ecdysteroids and juvenile hormones, which regulate development and reproduction. CYP expansion suggests that *G. sigillatus* may possess an enhanced ability to detoxify a broad range of harmful compounds, offering a selective advantage in chemically diverse environments and in energy conversion from agricultural by- products, thus contributing to more sustainable and economical cricket farming.

These findings highlight potentially distinctive genetics underlying adaptive traits in *G. sigillatus*, shedding light on its evolutionary trajectory within Orthoptera. The expansion of gene families related to chromatin remodelling, metabolism, digestion, and detoxification suggests that *G. sigillatus* has evolved genetic adaptations enhancing its physiological functions and ecological resilience, which is consistent with its global, cosmopolitan distribution in tropical regions and introduction into many non-native habitats. These adaptations likely support rapid responses to environmental stresses, optimise energy use, and strengthen defences against pathogens and toxins. Genomic resources will help to devise strategies for selectively breeding *G. sigillatus* for optimal reproductive traits in farming applications, as well as for unpicking the evolutionary causes and consequences of nuptial gift-giving, multiple mating, and mating system evolution in fundamental behavioural ecology research.

## Conclusion

There are over 6,000 species of crickets in the infraorder Gryllidea, over 60 of which are known to be consumed and, in some cases, farmed by humans (Cigliano et al., 2018; Magara et al., 2021). Generating a high-quality chromosome-scale assembly of one of the most globally widespread and commonly-farmed species, *Gryllodes sigillatus*, is a key step in promoting the adoption of genome-informed farming improvements for insects. The widespread farming of *G. sigillatus* is likely driven by their worldwide, cosmopolitan distribution and ease of collection (Weissman et al., 2012). However, there are many cricket species with farming potential, and the resources developed here will facilitate their study and use more broadly (Dossey et al., 2023; Magara et al., 2019, 2021). Recently, genomic resources, such as genome assemblies, transcriptome sequences, have increasingly deployed to improve strategies for farming crickets (Kataoka et al., 2022; Nakamura et al., 2022). Comparative and phylogenetic approaches continue to advance our understanding of the evolutionary history of crickets and allied taxa (Gray et al., 2020; Song et al., 2020; Dong et al., 2024), and genomic approaches in these species benefit from a longstanding history of major contributions to fundamental research in evolutionary and behavioural biology. Recent work, for example, has shed light on the genomics of speciation (Xu & Shaw, 2021; Yusuf et al., 2024), adaptation in the wild (Pascoal et al., 2020; Zhang et al., 2021; Zhang et al., 2024; Rayner et al. 2024), evolutionary developmental biology (Barnett et al., 2019; Ohde et al., 2022), and sperm competition, mate choice, and sexual selection (Simmons et al., 2013; Xu & Shaw, 2019). In addition, our findings provide valuable resources for precisely identifying genetic markers using approaches such as genome-wide association study (GWAS) and marker-assisted selection, and for guiding the genomic editing of breeding traits through RNA*i* or CRISPR-Cas9. These technologies have been successfully applied in other farmed crickets, including *Acheta domesticus* (Dossey et al., 2023) and *Gryllus bimaculatus* (Watanabe et al., 2017), and could be similarly implemented in *G. sigillatus*. Future research efforts will benefit not only from exploring the potential for *G. sigillatus* in greater depth using these genome-informed tools for applied and basic research, but also developing genomic resources across a broader taxonomic sampling of commonly studied Gryllids.

## Supplementary information

Supplementary Materials include Figs. S1, Table S1 to S7, Data table S1.

## Data availability

All genomic data (long-reads sequencing data, and Hi-C sequencing data) have been deposited at European Nucleotide Archive (ENA) under PRJEB84028. Transcriptome data have been deposited at PRJEB85996 and PRJNA1204288. Sequences of the genome assembly is publicly available at ENA under accession GCA_965111905. Annotation files of the genome assembly has been deposited at Zenodo as DOI: 10.5281/zenodo.14617131.

## Acknowledgements

The authors gratefully acknowledge funding support from the Australian Research Council to JH (DP180101708), the UK Natural Environment Research Council to NWB (NE/W001616/1), the China Scholarship Council to SZ (202106180022), and the United States National Science Foundation to SKS, BMS, and JH (IOS 16–54028). This research was also supported in part by the U.S. Department of Agriculture, Agricultural Research Service (Project Number: 6064- 32000-001-000D to KRD and 5010-22410-023-00-D to JLR). We thank the UK Biotechnology and Biological Sciences Research Council (BB/S019669/1 and BB/X019683/1) and James Hutton Institute for access to the UK’s Crop Diversity Bioinformatics High Performance Computing Cluster, with particularly acknowledgement of Ian Milne’s inexhaustible patience as sysadmin. Lianne Baker assisted with sample shipping and Tanya Sneddon with DNA extractions. The authors are grateful for constructive feedback on the manuscript from Xin Du. Mention of trade names or commercial products in this publication is solely for the purpose of providing specific information and does not imply recommendation or endorsement by the U.S. Department of Agriculture. USDA is an equal opportunity provider and employer.

## Author contributions

Conceived the study: JH, NWB. Designed experiments: all authors. Secured funding: JH, BMS, JLR, KRD, SKS, NWB. Performed experimental work: KRD, BF, SZ. Analysed data: KRD, BF, SZ. Led manuscript writing: SZ, NWB. Contributed to writing: JH, BF, SKS, BMS, JLR, KRD.

