## Supplementary information for "A high-quality reference genome and comparative genomics of the widely-farmed banded cricket (*Gryllodes sigillatus*) identifies selective breeding targets"

**This supplementary file includes:**

Figs. S1
Table S1 to S7

**Other Supplementary materials for this manuscript include:**

Supplementary data table 1

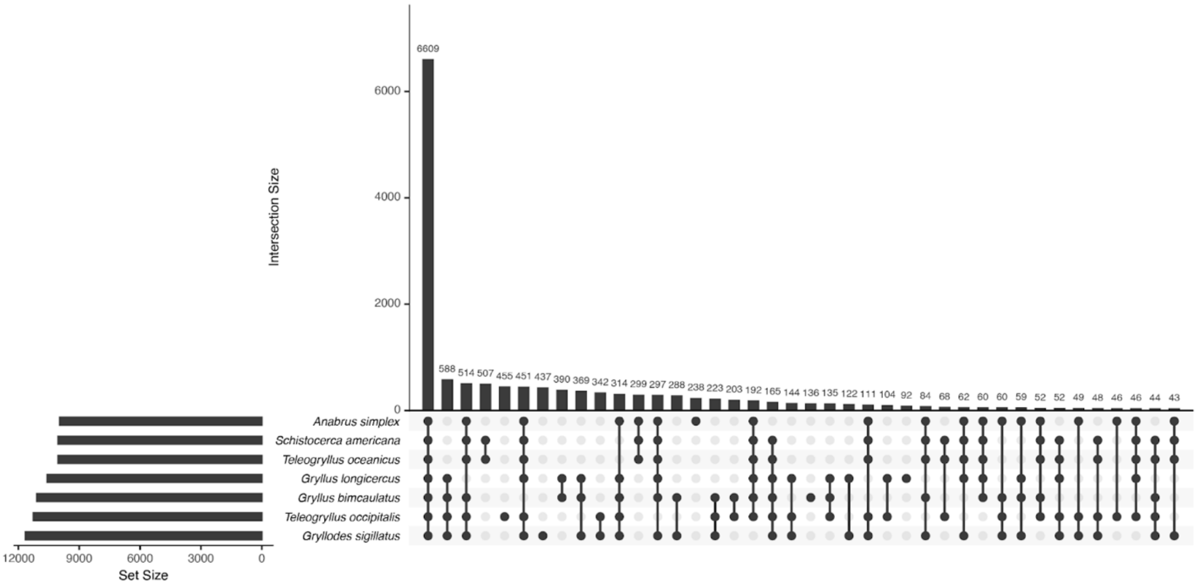


**Supplementary figure 1. UpSet plot illustrating the overlap of gene families among seven species.** The horizontal bars on the left represent the total number of gene families identified in each species. The vertical bars in the main panel indicate the number of gene families shared among the species combinations connected by the filled dots below.

**Supplementary table 1. Summary of the raw sequencing data.**

| Library Type | Base (Gb) | Average read length (bp) | Coverage Depth (×) |
|---|---|---|---|
| PacBio HiFi | 96.25 | 17,427.4 | 48.13 |
| Hi-C | 151.48 | 150 | 72.33 |

**Supplementary table 2. Information of RNA-seq data used for protein-coding gene prediction.**

| Sample name | Tissue | Sex | Mapping ratio (%) |
|---|---|---|---|
| F3AG10 | Accessory Glands | Male | 94.58 |
| F3AG2 | Accessory Glands | Male | 94.64 |
| F3AG8 | Accessory Glands | Male | 94.76 |
| F3H1 | Head | Female | 92.99 |
| F3H2 | Head | Female | 93.47 |
| F3H3 | Head | Female | 95.14 |
| F3RP1 | Ovaries | Female | 96.55 |
| F3RP2 | Ovaries | Female | 96.63 |
| F3RP3 | Ovaries | Female | 96.36 |
| KD1 | Whole body | Female | 94.81 |
| KD2 | Whole body | Female | 94.79 |
| KD3 | Whole body | Female | 94.58 |
| KD6 | Whole body | Male | 95.67 |
| KD7 | Whole body | Male | 96.78 |
| KD8 | Whole body | Male | 97.82 |

**Supplementary table 3. Statistics of *G. sigillatus* genome assembly.**

| Statistics | Value |
|---|---|
| **Total length (bp)** | 2,173,690,933 |
| **Contig number** | 455 |
| **Contig N50 (Mb)** | 9.02 |
| **Scaffold number** | 101 |
| **Scaffold N50 (Mb)** | 200.36 |
| **GC content (%)** | 39.77 |

**Supplementary table 4. Quality assessment of the assembled genome of *G. sigillatus*.**

| Chromosome name | Scaffold name | Contig number | Length (Mb) | QV | Error rate ($\times10^{-7}$ per base) |
|---|---|---|---|---|---|
| ChrX | scaffold_1 | 200 | 420.78 | 60.91 | 8.11 |
| Chr1 | scaffold_2 | 49 | 243.29 | 61.71 | 6.74 |
| Chr2 | scaffold_3 | 39 | 237.93 | 62.50 | 5.62 |
| Chr3 | scaffold_4 | 41 | 200.36 | 60.66 | 8.59 |
| Chr4 | scaffold_5 | 34 | 197.45 | 60.39 | 9.14 |
| Chr5 | scaffold_6 | 50 | 190.27 | 62.22 | 5.99 |
| Chr6 | scaffold_7 | 31 | 148.10 | 61.58 | 6.95 |
| Chr7 | scaffold_8 | 41 | 143.15 | 62.18 | 6.05 |
| Chr8 | scaffold_9 | 43 | 136.84 | 60.30 | 9.33 |
| Chr9 | scaffold_10 | 42 | 124.84 | 61.40 | 7.24 |
| Chr10 | scaffold_11 | 36 | 124.84 | 60.12 | 9.72 |
| **Entire genome** | | 455 | 2,173.70 | 61.21 | 7.56 |

**Supplementary table 5. Quality assessment of the assembled genome and gene set of *G. sigillatus* using BUSCOs.** The reference BUSCO set was insecta_odb10 with a total BUSCO number of 1367.

| **Object** | **Type** | **Number** | **Percent (%)** |
|---|---|---|---|
| Assembly | Complete BUSCOs (C) | 1360 | **99.5** |
| | Complete and single-copy BUSCOs (S) | 1342 | 98.2 |
| | Complete and duplicated BUSCOs (D) | 18 | 1.3 |
| | Fragmented BUSCOs (F) | 0 | 0 |
| | Missing BUSCOs (M) | 7 | 0.5 |
| Gene set | Complete BUSCOs (C) | 1347 | **98.5** |
| | Complete and single-copy BUSCOs (S) | 916 | 67.0 |
| | Complete and duplicated BUSCOs (D) | 431 | 31.5 |
| | Fragmented BUSCOs (F) | 4 | 0.3 |
| | Missing BUSCOs (M) | 16 | 1.2 |

**Supplementary table 6. Summary of repeat elements in *G. sigillatus* genome.**

| Repeats type | Number | Length (bp) | Percentage of Repeats (%) | Percentage of Genome (%) |
|---|---|---|---|---|
| **Transposable elements** | - | 996,365,344 | 94.90 | 45.77 |
| **SINEs** | 68,919 | 19,030,453 | 1.81 | 0.87 |
| **LINEs** | 400,691 | 253,317,115 | 24.13 | 12.64 |
| **LTRs** | 110,496 | 731,976 | 0.070 | 3.30 |
| **DNAs** | 413,510 | 170,341,526 | 16.22 | 7.82 |
| **Rolling circles** | 65,312 | 25,082,240 | 2.39 | 1.15 |
| **Unclassified** | 1,638,266 | 479,417,892 | 45.66 | 22.03 |
| **Small RNA** | 38,674 | 13,708,717 | 13.06 | 0.64 |
| **Satellites** | 613 | 173,884 | 0.017 | 0.01 |
| **Simple repeats** | 445,329 | 22,189,720 | 2.11 | 1.02 |
| **Low complexity** | 85,828 | 69,333 | - | 0.22 |
| **Total Repeats** | - | 1,049,877,180 | - | 48.22 |

**Supplementary table 7. Details of significantly expanded protein-coding gene families in *G. sigillatus*.** Gene families marked with an asterisk (*) represent those that failed to have a verified family name across Insecta but have homologous sequences in human gene families, indicating possible conserved functions across species.

| Ortholog identifier | Gene family name | Gene number in *G. sigillatus* | P value |
|---|---|---|---|
| OG0000001 | Histone H1/H5 | 52 | 2.88e-10 |
| OG0000005 | Trypsin | 23 | 0.0028 |
| OG0000009 | Cytochrome P450 | 21 | 0.040 |
| OG0000021 | Histone H2A | 26 | 1.16e-05 |
| OG0000022 | Histone H2B | 25 | 2.62e-05 |
| OG0000055 | *Unknown* | 20 | 1.8e-07 |
| OG0000060 | *Unknown* | 33 | 3.32e-15 |
| OG0000061 | *Unknown* | 16 | 0.00088 |
| OG0000065 | *Unknown* | 41 | 5.16e-23 |
| OG0000080 | *Unknown* | 17 | 8.25e-06 |
| OG0000104 | *Unknown* | 8 | 0.013 |
| OG0000106 | RTase | 9 | 0.0053 |
| OG0000117 | *Unknown* | 13 | 2.16e-05 |
| OG0000123 | LLRC* | 12 | 0.0044 |
| OG0000124 | THAP* | 15 | 1.47e-06 |
| OG0000125 | Complex I subunit 5 | 11 | 0.00029 |
| OG0000146 | Histone H3 | 27 | 2.22e-15 |
| OG0000197 | HARBI1* | 8 | 0.0018 |
| OG0000418 | *Unknown* | 10 | 0.00013 |
| OG0000450 | *Unknown* | 9 | 4.07e-05 |
| OG0000453 | *Unknown* | 12 | 1.36e-08 |
| OG0000494 | Complex I subunit 1 | 12 | 4.81e-07 |
| OG0000772 | *Unknown* | 5 | 0.012 |
| OG0000886 | SCM | 3 | 0.020 |
| OG0000941 | eIF-2B alpha/beta/delta subunits | 7 | 3.66e-05 |
| OG0001510 | Glutamate-gated ion channel | 7 | 3.66e-05 |
| OG0001843 | *Unknown* | 4 | 0.0042 |
| OG0007868 | *Unknown* | 3 | 0.020 |
